# Haplotype-aware sequence alignment to pangenome graphs

**DOI:** 10.1101/2023.11.15.566493

**Authors:** Ghanshyam Chandra, Daniel Gibney, Chirag Jain

## Abstract

Modern pangenome graphs are built using haplotype-resolved genome assemblies. During read mapping to a pangenome graph, prioritizing alignments that are consistent with the known haplotypes has been shown to improve genotyping accuracy. However, the existing rigorous formulations for sequence-to-graph co-linear chaining and alignment problems do not consider the haplotype paths in a pangenome graph. This often leads to spurious read alignments to those paths that are unlikely recombinations of the known haplotypes.

In this paper, we develop novel formulations and algorithms for haplotype-aware sequence alignment to an acyclic pangenome graph. We consider both sequence-to-graph chaining and sequence-to-graph alignment problems. Drawing inspiration from the commonly used models for genotype imputation, we assume that a query sequence is an imperfect mosaic of the reference haplotypes. Accordingly, we extend previous chaining and alignment formulations by introducing a recombination penalty for a haplotype switch. First, we solve haplotype-aware sequence-to-graph alignment in *O*(|*Q*| | *E*| |ℋ|) time, where *Q* is the query sequence, *E* is the set of edges, and ℋ is the set of haplotypes represented in the graph. To complement our solution, we prove that an algorithm significantly faster than *O*(|*Q*| | *E*| |ℋ|) is impossible under the Strong Exponential Time Hypothesis (SETH). Second, we propose a haplotype-aware chaining algorithm that runs in *O*(|ℋ| *N* log |ℋ|*N*) time after graph preprocessing, where *N* is the count of input anchors. We then establish that a chaining algorithm significantly faster than *O*(|ℋ|*N*) is impossible under SETH. As a proof-of-concept of our algorithmic solutions, we implemented the chaining algorithm in the Minichain aligner (https://github.com/at-cg/minichain). We demonstrate the advantage of the algorithm by aligning sequences sampled from human major histocompatibility complex (MHC) to a pangenome graph of 60 MHC haplotypes. The proposed algorithm offers better consistency with ground-truth recombinations when compared to a haplotype-agnostic algorithm.

## 1 Introduction

Pangenome graphs represent the maximum possible diversity that exists in the population [55]. A variety of methods have been developed that use pangenome graphs for common applications including genotyping, variant calling, haplotype reconstruction, etc. (see [10] for a review). Efforts toward using pangenome graphs have been further catalyzed by the advent of long and accurate reads. The recent progress in long-read technologies and assembly algorithms enables high-quality haplotype-phased assembly of human genomes [7,41,48,62]. Building a pangenome graph directly from phased assemblies instead of variant calls (VCF files) allows a more comprehensive representation of the variation [61]. Accordingly, the latest methods for pangenome graph construction use phased assemblies to construct a graph [9,19,15,31,28]. The input haplotypes are stored as paths in the graph.

Pangenome graphs raise fundamental questions about the trade-offs between complexity and usability. An arbitrary path in a pangenome graph corresponds to either the original haplotypes or their recombinations. The number of sequences spelled by a graph increases combinatorially with the number of variants. This issue has been addressed previously by using different techniques, e.g., by limiting the amount of variation in the graph [23,44,58], artificially simplifying complex regions [16], or restricting the alignment to either one [38,57,56] or two haplotype paths [4]. A more principled approach to tackle this problem may be to leverage the correlations between two or more genetic variants, i.e. where individuals tend to have same genotype [9,18]. For example, PanGenie is an alignment-free genotyping algorithm that leverages long-range haplotype information inherent in the phased genomes [9].

A pangenome graph is represented either as a directed cyclic graph or a directed acyclic graph (DAG), where each vertex is labeled by a sequence. The primary formulation for sequence-to-graph alignment problem seeks a path in the graph which spells a sequence with minimum edit distance from the query sequence [2,14,24,25,40,36,46]. *O*(|*V*| + |Q| |E|)-time algorithms exist for both exact and approximate pattern matching problems for graphs, where *Q* denotes the query sequence, *E* denotes the set of edges, and *V* denotes the set of vertices. These formulations do not consider the associations between genetic variants and may lead to alignments with spurious recombinations in variant-dense regions [44]. Co-linear chaining is another common technique used in modern aligners. It is used to identify a coherent subset of anchors (short exact matches) that can be joined together to produce an alignment. The existing formulations for chaining on graphs share the same limitation of not considering the associations between genetic variants [6,32,35,45,49]. Some of these chaining algorithms run in *O*(*KN* log *KN*) time after graph preprocessing [6,32], where *K* denotes the minimum number of paths required to cover all the vertices and *N* denotes the count of input anchors.

In this paper, we address the above limitations by introducing ‘haplotype-aware’ formulations for (i) sequence-to-DAG alignment and (ii) sequence-to-DAG chaining problems. Our formulations use the haplotype path information available in modern pangenome graphs. The formulations are inspired from the classic Li-Stephens haplotype copying model [30]. The Li-Stephens model is a probabilistic generative model which assumes that a sampled haplotype is an imperfect mosaic of known haplotypes. Similarly, our formulation for haplotype-aware sequence-to-DAG alignment problem optimizes the number of edits and haplotype switches simultaneously (Figure 1). We give *O*(|*Q*| | *E*| |ℋ|) time algorithm for solving the problem, where ℋ denotes the set of haplotypes represented in the pangenome DAG. We further prove that the existence of a significantly faster algorithm than *O*(|*Q*| | *E*| |ℋ|) is not possible under the strong exponential time hypothesis (SETH) [59]. We also formulate the haplotype-aware co-linear chaining problem. We solve it in *O*(|ℋ| *N* log |ℋ| *N*) time, assuming that a one-time preprocessing of the DAG is done in *O*(|*E*| |ℋ|) time. To provide evidence of the near optimality of this algorithm, we show that it is impossible to solve this problem significantly faster than *O*(|ℋ|*N*) under SETH.

**Fig. 1:**
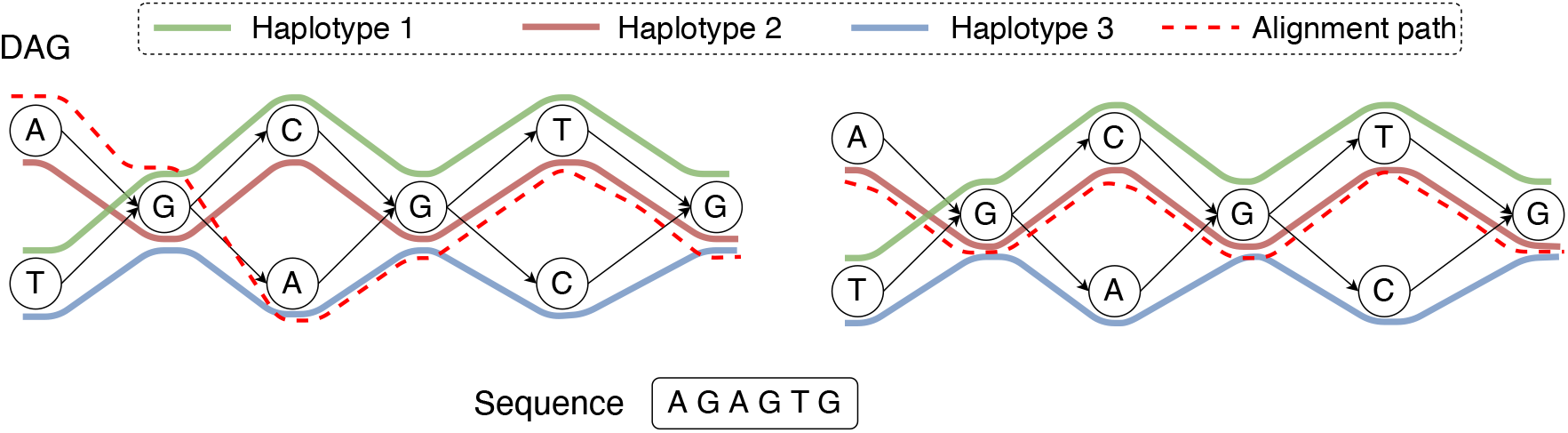
An example of an acyclic pangenome graph built using three haplotypes. The left figure shows the optimal alignment of the query sequence to the DAG with minimum edit distance. Accordingly, the edit distance is zero and the count of recombinations is two. Next, suppose that we use recombination penalty −5 (formally defined in Section 2). The right figure shows the new optimal alignment, where there is no recombination because the alignment path is consistent with Haplotype 2.

We implemented the proposed chaining algorithm in Minichain and evaluated it using simulated and real sequences. We built a pangenome DAG using 60 publicly-available complete major histocompatibility complex (MHC) sequences [27]. We simulated query MHC haplotypes as mosaics of the 60 haplotypes. We demonstrate that introducing recombination penalty in the formulation leads to a much better consistency between the observed recombination events and the ground truth. We achieved Pearson correlation up to 0.94 between the observed recombination count and the ground-truth recombination count, whereas the correlation remained below 0.32 if recombinations are not penalized. We also tested Minichain using publicly-available PacBio and Oxford Nanopore long reads from CHM13 human genome [41,60]. Here we demonstrate a better consistency between the ground-truth haplotype (CHM13) and the selected haplotype in the output. Our experiments suggest that haplotype-aware pattern matching to pangenome graphs can be useful to improve read mapping and variant calling accuracy.

## 2 Methods for haplotype-aware sequence-to-graph alignment

### 2.1 Technical background and notations

#### Haplotype-aware pangenome DAG

Let *G*(*V, E, σ, ℋ*) be a DAG representing a pangenome reference. We assume that the DAG is connected and |*E*| ≥ |*V* | − 1. Let *σ* denote an alphabet. Function *σ* : *V* → *σ* assigns a character label to each vertex. ℋ = {*H*_1_, *H*_2_, …, *H*_|ℋ|_} denotes the set of haplotype sequences represented in the DAG. Each haplotype sequence can be identified using a path in the DAG. We define *haps*(*v*) as {*i* | *H*_*i*_ includes vertex *v}*. We assume that *haps*(*v*) ≠ *ϕ* for all *v* ∈ *V* and *haps*(*u*) ∩ *haps*(*v*) ≠ *ϕ* for all edges (*u, v*) ∈ *E*. In other words, each vertex and edge must be supported by at least one haplotype. Given a query sequence *Q* ∈ *Σ*^+^, we will find its exact and approximate matches in the DAG. For brevity, we use [*m*] to denote the set {1, 2, …, *m*}, *m* ∈ ℕ. Let an *alignment path* of length *l* in the DAG be denoted as an ordered sequence (*w*_1_, *w*_2_, …, *w*_*l*_), where each *w*_*i*_ is a pair of the form (*v, h*) such that *v* ∈ *V, h* ∈ *haps*(*v*), and (*w*_*i*_.*v, w*_*i*+1_.*v*) ∈ *E* for all *i* ∈ [*l* − 1]. Thus, in our haplotype-aware setting, an alignment path specifies a path in the DAG and the indices of the selected haplotypes along the path. We say that a *recombination* has occurred in between *w*_*i*_ and *w*_*i*+1_ if *w*_*i*_.*h* ≠ *w*_*i*+1_.*h*. Given an alignment path 𝒫 in the DAG, we use functions *s*(𝒫) and *r*(𝒫) to denote the sequence spelled by the path and the count of recombinations in the path, respectively.

#### Hardness result for the Edit Distance problem

Our hardness result (Lemma 3) will build on top of the seminal work of Backurs and Indyk [5]. The *Orthogonal Vector (OV) Problem* asks: given two sets of *d*-dimensional binary vectors *A* and *B*, |*A*| = |*B*| =: *n*, decide if there is a pair *a* ∈ *A, b* ∈ *B* such that *a* · *b* = 0. If strong exponential time hypothesis (SETH) [59] holds and *d* = *ω*(log *n*), there is no *ϵ >* 0 for which an *O*(*n*^2−*ϵ*^poly(*d*)) algorithm can solve the OV problem [63]. This result is frequently used to study the hardness of other problems [5,13,14,17,20]. For any two sequences *P*_1_ and *P*_2_, let EDIT(*P*_1_, *P*_2_) be the edit distance between them. Let us define another distance measure between two sequences as following:

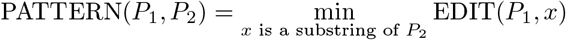

Backurs and Indyk [5] proved the hardness of computing edit distance by designing two reductions: (i) OV to PATTERN, and (ii) PATTERN to EDIT. We recall the properties of their first reduction gadget.

##### Lemma 1

**(ref. [5])**

*Given an OV instance with sets A* = {*a*_1_, …, *a*_*n*_} *and B* = {*b*_1_, …, *b*_*n*_} *comprising d-dimensional binary vectors, there exist transformations of sets A and B into two sequences S*_1_ *and S*_2_ *respectively over the alphabet* {0, 1, 2} *such that the following properties hold:*

– *Computing* PATTERN(*S*_1_, *S*_2_) *determines the existence of orthogonal vectors. There exist constants β*_1_, *β*_2_ ∈ ℕ *such that if there is a pair of orthogonal vectors a* ∈ *A, b* ∈ *B, then* PATTERN(*S*_1_, *S*_2_) ≤ *β*_1_*n* − *β*_2_; *otherwise*, PATTERN(*S*_1_, *S*_2_) = *β*_1_*n*.
– |*S*_1_| *is a function of n and d. Similarly*, |*S*_2_| *is a function of n and d. Both* |*S*_1_| *and* |*S*_2_|*are in O*(*n* ·poly(*d*)).
– *Time to compute sequences S*_1_ *and S*_2_ *from sets A and B is in O*(*n* · poly(*d*)).

We denote the injective function that generates the gadget sequence *S*_1_ from set *A* as *f*_1_. Similarly, injective function *f*_2_ generates sequence *S*_2_ from set *B*. The exact definitions of *f*_1_ and *f*_2_ are available in [5].

### 2.2 Problem definitions

The input to each of the following problem is a DAG *G*(*V, E, σ*,ℋ), a query *Q ∈ Σ*^+^, and a user-specified threshold *k* ∈ ℕ. In each problem, we seek a match of the query in the DAG while controlling the number of recombinations.

– *Problem 1 (Exact matching, limited recombinations):* Determine if there is an alignment path 𝒫 in DAG *G* such that *EDIT* (*s*(𝒫), *Q*) = 0 and *r*(𝒫) *< k*.
– *Problem 2 (Approximate matching, no recombination):* Determine if there is an alignment path 𝒫 in DAG *G* such that *EDIT* (*s*(𝒫), *Q*) *< k* and *r*(𝒫) = 0.
– *Problem 3 (Approximate matching, limited recombinations):* Let *k, c*_1_, *c*_2_ be user-specified thresholds, where *k* ∈ ℕ and *c*_1_, *c*_2_ ∈ ℝ_≥0_. Here *c*_1_ indicates cost per edit and *c*_2_ indicates cost per recombination. Determine if there is an alignment path 𝒫 in DAG *G* such that *c*_1_ *· EDIT* (*s*(𝒫), *Q*) + *c*_2_ *· r*(𝒫) *< k*.

In Problem 3, we assume a constant cost per recombination but one can also consider using different cost per locus based on known rates of recombination [43]. In the above set of problems, we have omitted the case with exact matching and no recombination. One way to solve this case is by doing exact sequence matching against each haplotype independently.

### 2.3 Proposed algorithms and complexity analysis

#### Lemma 2.

*Problems 1-3 can be solved in O*(|*Q*||*E*||ℋ|) *time*.

*Proof*. We show how to solve Problem 3 using dynamic programming. The same can be extended to Problems 1 and 2 as well. Our approach is similar to the standard sequence-to-DAG alignment algorithm [25] except that we also track recombinations. We add a dummy source vertex *v*_*ϵ*_ with an empty label. It has zero incoming edges and |*V*| outgoing edges to all *v* ∈ *V*. Assume *haps*(*v*_*ϵ*_) = [|ℋ|]. Let *D*(*i, v, h*) denote the minimum cost for aligning the (possibly empty) prefix of sequence *Q* ending at index *i* against an alignment path that ends at pair (*v, h*), *v* ∈ *V* ∪ {*v*_*ϵ*_} *, h* ∈ *haps*(*v*). We also maintain a second table *D*^*′′*^ such that *D*^*′′*^(*i, v*) will store min_*h*∈*haps*(*v*)_ *D*(*i, v, h*). Initialize *D*(0, *v, h*) = *D*^*′′*^(0, *v*) = 0 for all *v* ∈ *V*∪{*v*_*ϵ*_}, *h* ∈ *haps*(*v*). Initialize the remaining cells in table *D*^*′′*^ to ∞. For *i* ≥ 1, *v* ∈ *V* ∪ {*v*_*ϵ*_}, *h* ∈ *haps*(*v*), update *D*(*i, v, h*) and *D*^*′′*^(*i, v*) using the following recursion:

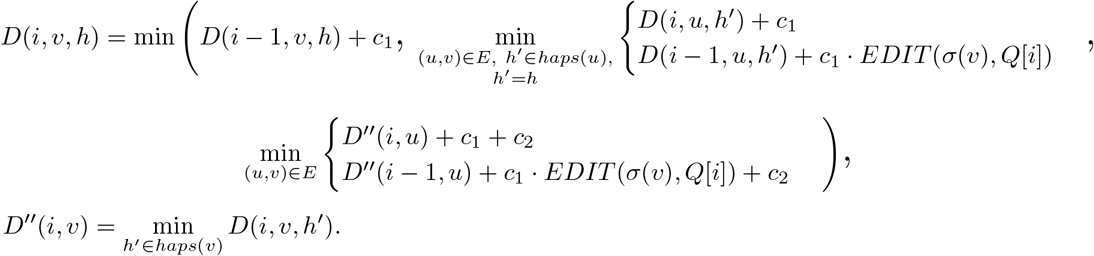

The recursion computes the optimal value of *D*(*i, v, h*) by considering the possibility of character deletion, insertion, match, and mismatch. The product *c*_1_ · *EDIT* (*σ*(*v*), *Q*[*i*]) equals zero if there is a match between characters *σ*(*v*) and *Q*[*i*]. It equals *c*_1_ in case of a mismatch. The other expressions involve addition of *c*_1_ to consider the possibility of an insertion or a deletion. The expressions involving the addition of *c*_2_ evaluate the possibility of a recombination. After updating *D*(*i, v, h*) for all *h* ∈ *haps*(*v*), we update *D*^*′′*^(*i, v*) using the above equation. We compute values in table *D* in the increasing order of *i*, and then the increasing topological order of *v* ∈ *V* ∪ {*v*_*ϵ*_} for a fixed *i*. The output, i.e., the minimum alignment cost is 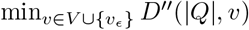. The runtime of the algorithm is dominated by the recursion. Solving the recursion takes 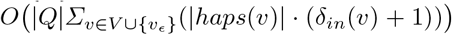 time, where *δ*_*in*_(*v*) denotes the in-degree of vertex *v*. As a result, the time complexity of the algorithm is *O*(|*Q*| | *E*| |ℋ|).

Next, we prove that a significantly faster algorithm than *O*(|*Q*| | *E*| |ℋ|) for solving Problem 2 or 3 is unlikely. The existence of a faster algorithm for Problem 1 remains open.

#### Lemma 3.

*For any constant ϵ >* 0, *there is no algorithm that solves Problem 2 or Problem 3 in O*(|*Q*|^1−*ϵ*^|*E*||*ℋ*| +|*Q*||*E*|^1−*ϵ*^|*ℋ*| + |*Q*||*E*||*ℋ*|^1−*ϵ*^) *time, unless SETH is false*.

*Proof*. Problem 3 is generalized version of Problem 2. Therefore, it suffices to prove the hardness of Problem 2. We derive the reduction from OV problem. Suppose we have an OV instance with equal sized sets *A* and *B* comprising -dimensional binary vectors. Define .*n* = |*A*| = |*B*|. We assume w.l.o.g. that *n* is a perfect square. We partition set *A* into 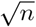 subsets 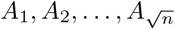, each containing 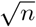 vectors. Similarly, we partition set *B* into 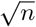 subsets 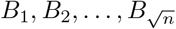, each containing 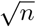 vectors. Observe that solving the OV problem for all (*A*_*i*_, *B*_*j*_) instances, 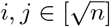, solves the OV problem for instance (*A, B*).

Next, we describe the construction of our gadget comprising query sequences 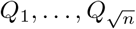 and haplotype-aware pangenome DAG *G*(*V, E, σ, ℋ*). Assume alphabet {0, 1, 2} for defining the query sequences and the vertex labels in the DAG. Define *Q*_*i*_ = *f*_1_(*A*_*i*_) for all 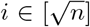 (ref. Lemma 1). Next, we will construct 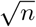 haplotype sequences. Define *H*_*i*_ = *f*_2_(*B*_*i*_) for all 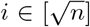. Using Lemma 1, we know that all our haplotype sequences have uniform length, i.e., 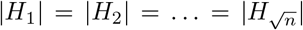. Next, we construct a DAG where these haplotype sequences can be represented. We build a DAG with |*H*_*i*_| ‘layers’. Each layer comprises three vertices labelled with characters ‘0’, ‘1’ and ‘2’ respectively (see Figure 2a). Each layer (except the last) is *fully connected* to its next adjacent layer using 3^2^ = 9 directed edges. Subsequently, we identify the unique path that spells each of the 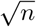 haplotype sequences in this DAG. Finally, we discard the vertices and edges that are not used by any haplotype sequence (Figure 2b).

**Fig. 2:**
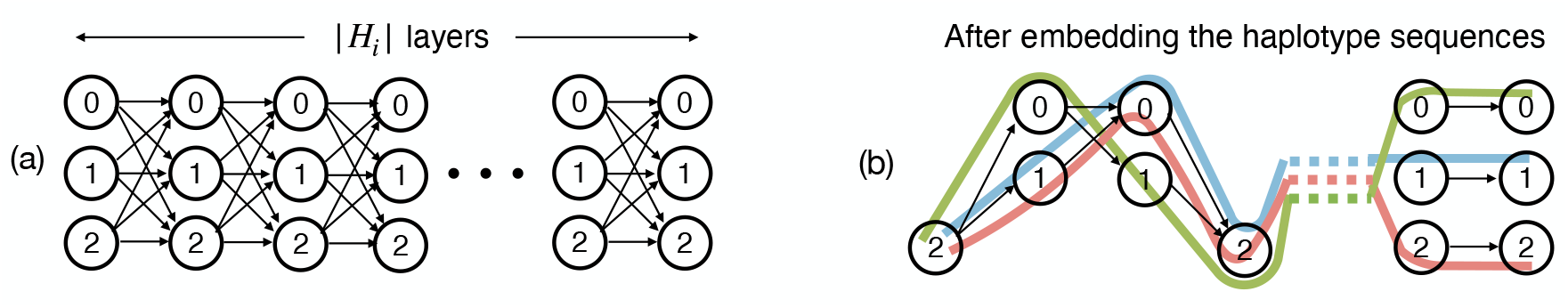
An illustration of the pangenome DAG used in the reduction proof of Lemma 3.

Next, the above gadget will allow us to solve the OV problem by invoking the haplotype-aware sequence to graph alignment algorithm 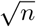 times, once for each of the 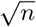 query sequences. An alignment path with no recombination in the DAG spells a substring of a haplotype. Using Lemma 1, we make the following claim. There exist two orthogonal vectors *a* ∈ *A* and *b* ∈ *B* if and only if we have at least one query sequence 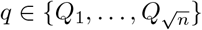 for which an alignment path *𝒫* exists such that *r*(𝒫) = 0 and *EDIT* 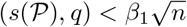. It is only remaining to show that a faster algorithm for Problem 2 will contradict SETH by enabling a faster algorithm for the OV problem. Observe that (i) each |*Q*_*i*_| and |*H*_*i*_| is in 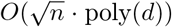, (ii) |*V* | and |*E*| are asymptotically equal to |*H*_*i*_|, (iii) number of haplotypes is 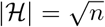, and (iv) we invoke the alignment algorithm 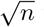 times. Construction of the gadget requires *O*(*n·* poly(*d*)) time. Therefore, if Problem 2 can be solved in *O*(|*Q*|^1−*ϵ*^|*E*||*ℋ*| + |*Q*||*E*|^1−*ϵ*^|*ℋ*| + |*Q*||*E*||*ℋ*|^1−*ϵ*^) time for some *ϵ >* 0, then the OV problem can be solved in *O*(*n*^2−*ϵ/*2^ *·* poly(*d*)) time.

## 3 Methods for haplotype-aware chaining on graphs

Most alignment tools use seed-chain-extend heuristic to compute the alignments quickly [53,54]. Given a set of seed matches (anchors) as input, co-linear chaining is a rigorous optimization technique to identify promising alignment regions in a reference. It identifies a coherent subset of anchors such that their coordinates are ordered on the query and the reference. The selected anchors are subsequently combined together to form an alignment. Several versions of co-linear chaining problems have been studied for aligning two sequences [1,22,34,39,11,12,42]. Recent works have further studied the extension of chaining on acyclic [35,32,6,49] and cyclic pangenome graphs [3,45] but these formulations do not consider the haplotype paths.

### 3.1 Technical background and notations

From hereon we assume that vertices of a haplotype-aware pangenome DAG are labeled with sequences. Sequence labeled vertices permit a more compact graph representation which will be useful in the context of chaining. It is trivial to transform a graph from sequence-labeled form to character-labeled form, and vice versa. Let function *σ* : *V*→ *Σ*^+^ label each vertex *v* ∈ *V* with sequence *σ*(*v*) in DAG *G*(*V, E, σ*, ℋ). The input to the chaining problem is a set of anchors {*M* [1], *M* [2], …, *M* [*N*]}. An anchor is represented by a 3-tuple (*v*, [*x*..*y*], [*c*..*d*]), which signifies a match between query substring *Q*[*c*..*d*] and substring *σ*(*v*)[*x*..*y*] in the DAG (Figure 3). The *weight* function assigns a user-specified weight to each anchor. Anchor weights are typically set proportional to the length of the matching substrings in practice. We use *R*^−^(*v*) ⊆ *V* to denote the subset of vertices that can reach *v* using a path in the DAG. Vertex *v* is also included in *R*^−^(*v*). Next, we define the precedence relationship between a pair of anchors.

**Fig. 3:**
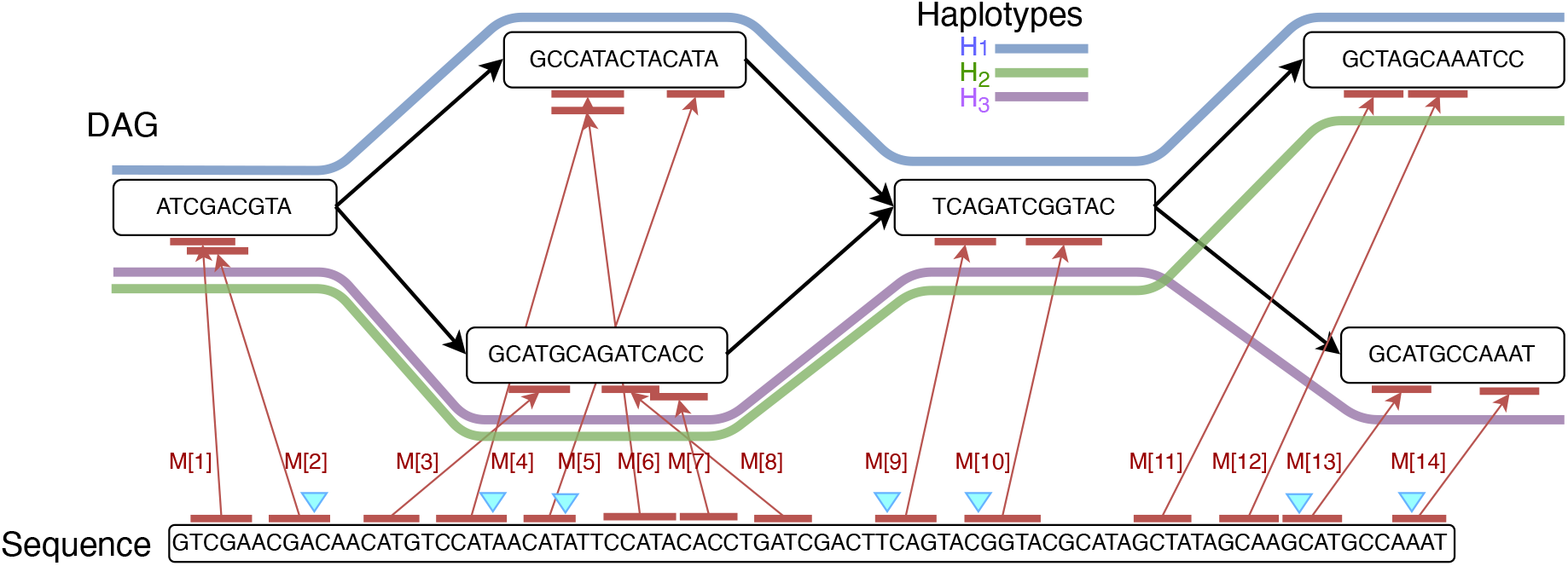
An example that shows anchors (in red) between a query sequence and a pangenome DAG. The DAG comprises three haplotype sequences *H*_1_, *H*_2_, *H*_3_. In this example, the ordered sequence of pairs ((2, 1), (4, 1), (5, 1), (9, 1), (10, 1), (13, 3), (14, 3)) forms a chain and the corresponding anchors are highlighted using blue markers. This chain includes a single recombination.

#### Definition 1 (Precedence)

*Given two anchors M* [*i*] *and M* [*j*], *M* [*i*] *precedes (*≺*) M* [*j*] *if (i) M* [*i*].*d < M* [*j*].*c, (ii) M* [*i*].*v* ∈ *R*^−^(*M* [*j*].*v*), *and (iii) M* [*i*].*y < M* [*j*].*x if M* [*i*].*v* = *M* [*j*].*v*.

#### Definition 2 (Chain)

*A chain is an ordered sequence* (*s*_1_, *s*_2_, …, *s*_*p*_), *where each s*_*j*_ *is a pair of the form* (*i, h*) *such that i* ∈ [*N*], *h* ∈ *haps*(*M* [*i*].*v*) *and M* [*s*_*j*_.*i*] ≺ *M* [*s*_*j*+1_.*i*] *for all j* ∈ [*p* − 1].

In our definition, a chain specifies the indices of the selected haplotypes alongside the anchors (Figure 3). We say that a recombination has occurred in between *s*_*j*_ and *s*_*j*+1_ if *s*_*j*_.*h* ≠ *s*_*j*+1_.*h*. Given a chain *S*, we use function *r*(*S*) to denote the count of recombinations in *S*. Let *γ* ∈ ℝ_≥0_ be a user-specified parameter that specifies cost per recombination.

Similar to [35], we will use *path cover* to facilitate efficient sparse dynamic programming on the DAG. A path cover is a set of paths in the DAG such that every vertex in *V* is included in at least one path. Previous chaining algorithms, e.g., in [6,32,35], identify a path cover with the minimum number of paths because the runtime of sparse dynamic programming increases with the size of the path cover [35]. In the haplotype-aware setting, we will directly use the paths corresponding to the haplotype sequences as path cover. Let *P*_1_, *P*_2_, …, *P*_|*ℋ*|_ be the paths corresponding to the haplotype sequences *H*_1_, *H*_2_, …, *H*_|*ℋ*|_ respectively. {*P*_1_, *P*_2_, …, *P*_|*ℋ*|_} is a path cover of the DAG because each vertex in the DAG is included in at least one haplotype (Section 2.1). Suppose *last*2*reach*(*v, h*), *v* ∈ *V, h* ∈ [|*ℋ*|] denotes the last vertex in path *P*_*h*_ that belongs to *R*^−^(*v*). *last*2*reach*(*v, h*) does not exist if no vertex in path *P*_*h*_ can reach vertex *v*. We precompute *last*2*reach*(*v, h*) for all *v* ∈ *V* and *h* [|ℋ|] in *O*(|*E*| |ℋ|) time by using the algorithm from Makinen *et al*. [35]. This is a one-time preprocessing step that will be useful to solve the chaining problems efficiently. Next, we also use the following standard data structure in our chaining algorithm.

#### Lemma 4

**(ref. [33])**

*Let n be the number of leaves in a tree, each storing a* (*key, value*) *pair. The data structure uses a balanced binary search tree. It supports update and RMQ (range maximum query) operations, defined below, in O*(log *n*) *time:*

– *update*(*k, val*): *For the leaf w with key* = *k, value*(*w*) ← max(*value*(*w*), *val*).
– *RMQ*(*l, r*): *Return* max {*value*(*w*) | *l < key*(*w*) *< r*}.

*Given n* (*key, value*) *pairs, the tree can be constructed in O*(*n* log *n*) *time and O*(*n*) *space*.

### 3.2 Problem definitions

*– Problem 4 (Limited recombinations):* Determine an optimal chain *S* = *s*_1_, *s*_2_, …, *s*_*p*_ that maximises the chaining score, defined as 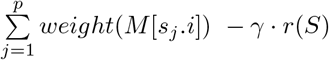.

The second term *γ*· *r*(*S*) in the above scoring function corresponds to the penalty for haplotype switches in a chain. While scoring a chain, it is also essential to penalise the gap corresponding to the distance (in basepairs) between adjacent pair of anchors. Accordingly, we formulate a gap-sensitive version of the above problem. We assume the same definition of function *gap*(*M* [*i*], *M* [*j*]), *i, j* ∈ [*N*] as used in [6, Problem 2a].

*– Problem 5 (Limited recombinations, gap-sensitive):* Determine an optimal chain *S* = *s*_1_, *s*_2_, …, *s*_*p*_ that maximises the chaining score, defined as 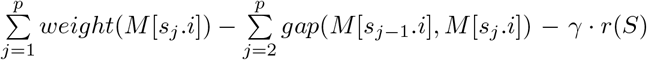.

### 3.3 Proposed algorithms and complexity analysis

First, it is useful to discuss a simple dynamic programming solution for Problem 4. Let *C*(*i, h*) denote the optimal score of a chain ending at pair (*i, h*), where *i* ∈ [*N*], *h* ∈ *haps*(*M* [*i*].*v*). We use *C*^*′′*^(*i*) to store *max*_*h*∈*haps*(*M*[*i*].*v*)_*C*(*i, h*). A single anchor is a valid chain of size one; therefore, we initialize *C*(*i, h*) to *weight*(*M* [*i*]) for all *i* ∈ [*N*], *h* ∈ *haps*(*M* [*i*].*v*). Also, initialize *C*^*′′*^(*i*) to *weight*(*M* [*i*]) for all *i* ∈ [*N*]. We update *C*(*i, h*) to its optimal value using the following recursion:

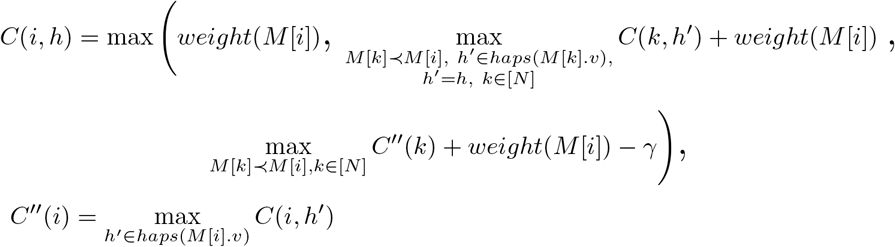

If *M* [*k*] ≺ *M* [*i*] and path *P*_*h*_ covers both the vertices *M* [*k*].*v* and *M* [*i*].*v*, then the chaining score obtained using *C*(*k, h*) is considered without recombination penalty. The third term in the expression considers the score with recombination penalty *γ*. This is needed when (*k, h*^*′*^), *h*^*′*^ ≠ *h* precedes (*i, h*). After updating *C*(*i, h*) for all *h* ∈ *haps*(*M* [*i*].*v*), we also update *C*^*′′*^(*i*) using the above equation. Let function *rank*(*v*) specify the topological ordering of vertex *v* in the DAG. The *C*(*i, h*) values should be computed in lexicographically ascending order based on the key (*rank*(*M* [*i*].*v*), *M* [*i*].*x*). For a fixed *i, C*(*i, h*) values may be computed in any order. The above algorithm fills up to |ℋ|*N* values in table *C*. Computing each value using a linear scan would require considering all *N* number of anchors and filtering those which satisfy the precedence criteria. Even if we ignore the time for filtering and checking precedence, the worst-case asymptotic runtime of this algorithm grows at least as fast as |ℋ| *N* ^2^. This makes it too slow when *N* is in millions, which typically happens when aligning megabase-long *de novo* assembled sequences to human pangenome DAGs. To address this, we propose a faster *O*(|ℋ| *N* log|ℋ| *N*) time algorithm for solving Problem 4. First, we note the following inequality.

#### Lemma 5.

*C*(*i, h*) ≥ *C*^*′′*^(*i*) − *γ for all i* ∈ [*N*], *h* ∈ *haps*(*M* [*i*].*v*).

*Proof*. Consider any value of *i* ∈ [*N*] and *h* ∈ *haps*(*M* [*i*].*v*). Let *h*^*′*^ = argmax_*k haps*(*M* [*i*].*v*)_ *C*(*i, k*), breaking ties arbitrarily if necessary. Then, *C*^*′′*^(*i*) equals *C*(*i, h*^*′*^). If *h*^*′*^ = *h*, then *C*(*i, h*) = *C*^*′′*^(*i*), thus the inequality holds. Next, consider the case when *h*^*′*^ ≠ *h*. Let *S*_1_ = *s*_1_, *s*_2_, …, *s*_*p*−1_, *s*_*p*_ be an optimal chain that ends at pair (*i, h*^*′*^). We consider another chain 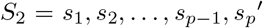 such that 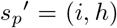. Suppose the score of chain *S*_2_ is *x*. Both chains *S*_1_ and *S*_2_ differ only by the haplotype used in their last pairs. Therefore, the difference in the chaining scores of *S*_1_ and *S*_2_ is at most *γ* (one recombination). Thus, *x* ≥ *C*^*′′*^(*i*) − *γ*. Also, *C*(*i, h*) must be ≥ *x*.

#### Lemma 6.

*Assuming DAG G*(*V, E, σ, ℋ*) *is preprocessed, Problem 4 can be solved in O*(|*ℋ*|*N* log |*ℋ*|*N*)*time*.

*Proof*. See Algorithm 1 for an outline of our approach. We extend the sparse dynamic programming framework of Makinen *et al*. [35] to our haplotype-aware setting. Our path cover {*P*_1_, *P*_2_, …, *P*_|*ℋ*|_} contains |*ℋ*| paths corresponding to the haplotype sequences. We use search trees 𝒯_1_, 𝒯_2_, …, 𝒯 _|*ℋ*|_, one per path. Each search tree 𝒯_*h*_ will record *C*(*i, h*) scores specific to that haplotype.

We initialise each search tree with keys {*M* [*i*].*d* 1 ≤ *i* ≤ *N*} and − ∞values (Line 1). In Lines 4-13, we create an array *Z* with *O*(|ℋ|*N*) tuples. After sorting of *Z* (Line 15), these tuples ensure that the search trees are updated and queried in a proper order. At the time of processing *C*(*i, h*), *i* ∈ [*N*], *h* ∈ *haps*(*M* [*i*].*v*), the algorithm would have already finished processing *C*(*k, h*^*′*^) for all *k, h*^*′*^ such that *M* [*k*] *M* [*i*], *h*^*′*^ ∈ *haps*(*M* [*k*].*v*). The score *C*(*k, h*^*′*^) would have already been recorded in search tree 𝒯_*h*_^*′*^ (Line 26).

All the anchors that lack a preceding anchor are optimally processed at the initialization stage (Line 2). Suppose an optimal chain ending at (*i, h*) is *s*_1_, *s*_2_, …, *s*_*p*−1_, *s*_*p*_ such that *s*_*p*−1_ = (*k, h*^*′*^) and *s*_*p*_ = (*i, h*). Assume that *C*(*k, h*^*′*^) is already computed optimally. While calculating *C*(*i, h*), the algorithm handles the following three cases:

– (Case 1) *h* = *h*^*′*^: In this case, we optimally compute *C*(*i, h*) by executing a range query in search tree *T*_*h*_ (Line 19). Since *M* [*k*] ≺ *M* [*i*] and *h*^*′*^ = *h*, search tree *T*_*h*_ returns the value *C*(*k, h*^*′*^) stored in key *M* [*k*].*d*.
– (Case 2) *h* ≠ *h*^*′*^ and path *P*_*h*_^*′*^ does not cover *M* [*i*].*v*: Observe that *last*2*reach*(*M* [*i*].*v, h*^*′*^) must exist in this case because *M* [*k*].*v* must reach *M* [*i*].*v*. In Lines 10-12, we add tuples in array *Z* to ensure that *C*_*l*2*r*_(*i*) is updated by using a range query in search tree 𝒯_*h*_^*′*^ (Line 22). We also put a penalty *γ* due to recombination in this case. This update to *C*_*l*2*r*_(*i*) happens after all anchors in vertex *last*2*reach*(*M* [*i*].*v, h*^*′*^) have been processed and search tree 𝒯_*h*_^*′*^ has been updated because of our definition and ordering of the tuples in array *Z* (Lines 12, 15). Later when we update *C*(*i, h*) in Line 19, it gets its optimal value using *C*_*l*2*r*_(*i*).
– (Case 3) *h* ≠ *h*^*′*^ and path *P*_*h*_^*′*^ covers *M* [*i*].*v*: If (*s*_1_, *s*_2_, …, (*k, h*^*′*^), (*i, h*)) is an optimal chain ending at (*i, h*), then *s*_1_, *s*_2_, …, (*k, h*^*′*^), (*i, h*^*′*^) must be an optimal chain that ends at (*i, h*^*′*^). Accordingly, *C*(*i, h*^*′*^) = *C*(*i, h*) + *γ*. It implies *C*^*′′*^(*i*) ≥ *C*(*i, h*) + *γ*. Using the inequality in Lemma 5, we get *C*(*i, h*) = *C*^*′′*^(*i*) − *γ*. Accordingly, *C*(*i, h*) is updated to its optimal value in Line 24.

The total runtime of the algorithm is dominated by sorting of array *Z*. Sorting an array of size *O*(|*ℋ*|*N*) takes *O*(|*ℋ*|*N* log |*ℋ*|*N*) time. Initializing |*ℋ*| search trees takes *O*(|*ℋ*|*N* log *N*) time. The algorithm uses *O*(|*ℋ*|*N*) number of *O*(log *N*)-time RMQ and update operations on the search trees (Lemma 4).

In the above algorithm, the optimal chain can be obtained easily by adding backtracking pointers. We next show that any algorithm to solve Problem 4 has a time complexity that is not polynomially faster than *O*(|*ℋ*|*N*) under SETH.

#### Lemma 7.

*For any constant ϵ >* 0, *there is no algorithm that solves Problem 4 in O*(|ℋ| ^1−*ϵ*^*N* + |ℋ| *N* ^1−*ϵ*^) *time, unless SETH is false*.

#### Algorithm 1

*O*(|*ℋ*|*N* log |*ℋ*|*N*) time chaining algorithm for Problem 4

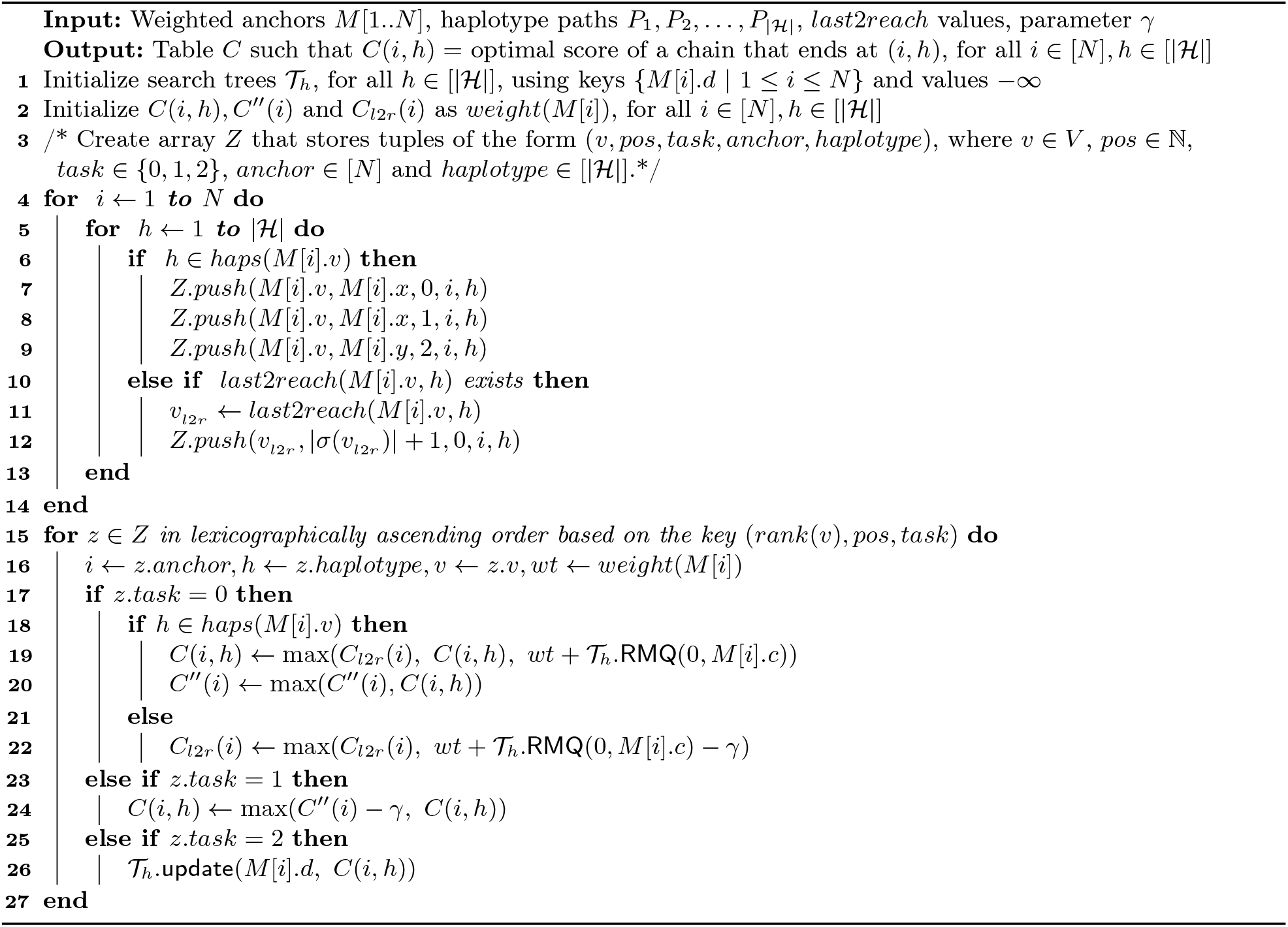

*Proof*. The reduction from the OV problem is as follows. Suppose an OV instance is given as *A* = {*a*_1_, …, *a*_*n*_} and *B* = {*b*_1_, …, *b*_*n*_} and each set is comprised of *d*-dimensional binary vectors. We first define component gadgets:

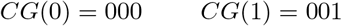

For two sequences *s*_1_ and *s*_2_, we use *s*_1_ ○ *s*_2_ or *s*_1_ *s*_2_ to denote concatenation of *s*_1_ and *s*_2_. For a symbol *c* and an integer *i*, we use *c*^*i*^ to denote symbol *c* repeated *i* times. Using the above component gadgets, we define our vector gadgets:

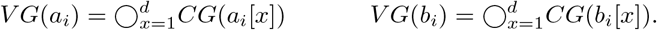

The query sequence is defined as 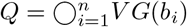. For 1 ≤ *i* ≤ *n*, we construct haplotype sequence *H*_*i*_ from vector *a*_*i*_:

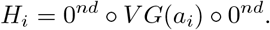

To create the vertices of graph *G* we start with: a vertex denoted *v*_*left*_ with label 0^*nd*^; 2*d* vertices denoted *v*_*x*,0_, *v*_*x*,1_ for 1 ≤ *x* ≤ *d* where each *v*_*x*,0_ has label *CG*(0) and each *v*_*x*,1_ has label *CG*(1); and a vertex denoted *v*_*right*_ with label 0^*nd*^. For edges, we add edges from *v*_*left*_ to *v*_1,0_ and from *v*_*left*_ to *v*_1,1_; for 1≤ *x < d*, we add edges from *v*_*x*,0_ to *v*_*x*+1,0_ and *v*_*x*+1,1_, and from *v*_*x*,1_ to *v*_*x*+1,0_ and *v*_*x*+1,1_; we then add edges from *v*_*d*,0_ to *v*_*right*_ and from *v*_*d*,1_ to *v*_*right*_. Next, we embed each haplotype in *G* by following its only possible path from *v*_*left*_ to *v*_*right*_ and deleting any vertices and edges not supported by some haplotype. See Figure 4. We call the subgraph of *G* excluding only *v*_*left*_ and *v*_*right*_ the *center* of *G*.

**Fig. 4:**
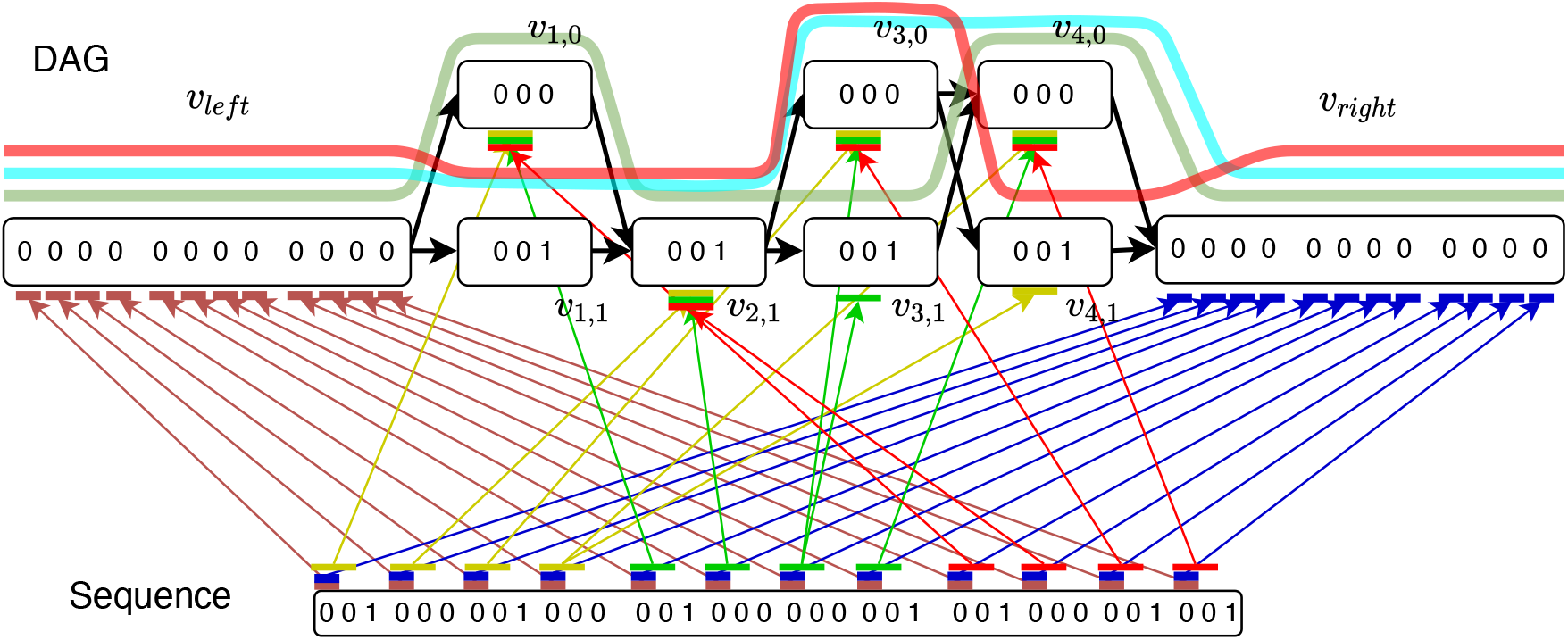
Reduction from OV problem with sets *A* = {0110, 1100, 1101} and *B* = {1010, 1001, 1011}.

The anchors are described next. For all *i* ∈ [*n*] and for all *x* ∈ [*d*], we define the following weight 1 anchors:

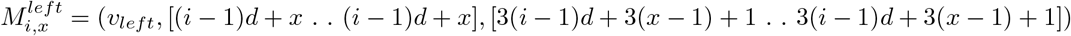

We similarly define

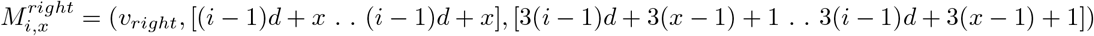

For all *i* ∈ [*n*] and for all *x* ∈ [*d*], if *b*_*i*_[*x*] = 0 and if *v*_*x*,0_ and *v*_*x*,1_ exist, we define the following weight 2 anchors

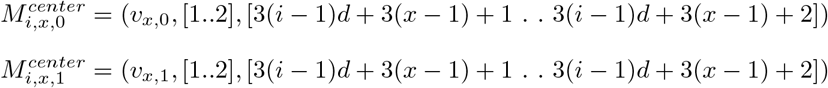

If either *v*_*x*,0_ or *v*_*x*,1_ do not exist, we omit the corresponding anchor. If *b*_*i*_[*x*] = 1 and *v*_*x*,0_ exists, we define the following weight 2 anchor

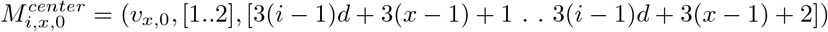

We set the recombination penalty *γ* = 1. We show the correctness of the above reduction in Lemmas 8-11.

#### Lemma 8.

*If there exists an orthogonal pair a*_*i*_ ∈ *A and b*_*j*_ ∈ *B, then there exists an anchor chain with cost* 2*d* + (*n* − 1)*d*.

*Proof*. First take anchors 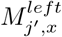 for 1 ≤ *j*^*′*^ *< j*, 1 ≤ *x* ≤ *d*. This part of the chain has a value of (*j* − 1)*d*. Next, for 1 ≤ *x* ≤ *d*, if *a*_*i*_[*x*] = 0 and *b*_*j*_[*x*] = 0 or *a*_*i*_[*x*] = 0 and *b*_*j*_[*x*] = 1 take 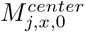; if *a*_*i*_[*x*] = 1 and *b*_*j*_ [*x*] = 1 take 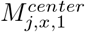. This part of the chain has a value of 2*d*. Lastly, for *j*^*′*^ *> j*, 1 ≤ *x* ≤ *d*, take anchors 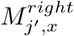, this part of the chain has value (*n* − *j*)*d*. This forms a valid chain and requires no recombinations. Thus, the total score is (*j* − 1)*d* + 2*d* + (*n* − *j*)*d* = 2*d* + (*n* − 1)*d*.

#### Lemma 9.

*If a chain exists with score* 2*d* + (*n* − 1)*d, then in this chain, no recombinations are used, and anchors starting at all the anchor starting positions in Q are used*.

*Proof*. For a given *x*, only one anchor to either *v*_*x*,0_ or *v*_*x*,1_ can be used while maintaining the precedence condition. Since each anchor to the center of *G* has weight 2, the total contribution of the center of *G* to the score is at most 2*d*. The remaining anchors in the chain have a weight of 1, and there are at most (*n* − 1)*d* of them after *d* anchors are used in the center. Hence, the maximum possible score is 2*d* + (*n*− 1)*d*.

The claim that there are no recombinations follows, as the maximum possible score using recombination becomes 2*d* + (*n* − 1)*d* − 1. The second claim follows since any chain using fewer than *dn* anchors can not achieve the maximum score of 2*d* + (*n* − 1)*d*.

#### Lemma 10.

*If a chain exists with the score* 2*d* + (*n* − 1)*d, then in this chain, all anchors for some vector gadget in Q are to the center of G*.

*Proof*. If anchors from two different vector gadgets in *Q* are to the center of *G*, then since the center is contributing 2*d* to the score (the only way a score of 2*d* + (*n* − 1)*d* is achieved), there must be anchors *M* and *M*^*′*^ from two different vector gadgets that occur in adjacent vertices in the center, say with *M* ≺ *M*^*′*^. However, this implies that there are at least *d* − 1 anchor positions in *Q* between *M* and *M*^*′*^ are not used in the solution, contradicting Lemma 9.

#### Lemma 11.

*If there exists a chain with score* 2*d* + (*n* − 1)*d, then there exists orthogonal vectors a*_*i*_ *and b*_*j*_.

*Proof*. By Lemma 10, some vector gadget *V G*(*b*_*j*_) has all of its anchors going to the center of *G*. By Lemma 9, no recombinations are used, and these must lie on some path for a haplotype constructed from a vector *a*_*i*_. By design of the component gadgets, for 1 ≤ *x* ≤ *d* if *a*_*i*_[*x*] = 1 we get anchor between *v*_*x*,1_ on the haplotype path for *h*_*i*_ and *V G*(*b*_*j*_) if and only if *b*_*j*_[*x*] = 0. We conclude that *a*_*i*_ and *b*_*j*_ are orthogonal.

Observe that the total number of *M*^*left*^ anchors is *dn*, the total number of *M*^*right*^ anchors is *dn*, and the total number of *M*^*center*^ anchors is at most 2*dn*, thus *N* = *O*(*dn*). The number of haplotypes is |ℋ|= *n*. Furthermore, *Q* = *O*(*dn*) and the graph *G* has *O*(*d*) vertices and edges, and the sum of vertex label lengths is *O*(*dn*). Hence, the reduction takes *O*(*dn*) time. Combined with Lemmas 8 and 11, it follows that an algorithm for Problem 4 running in time *O*(|ℋ|^1−*ϵ*^*N* + |ℋ| *N* ^1−*ϵ*^), solves the OV problem in time *O*(*n*^2−*ϵ*^ poly(*d*)). This completes the proof of Lemma 7.

Problem 5 can be solved by extending Algorithm 1 (Supplementary Document). The extension requires a few additional terms corresponding to gap cost between adjacent anchors. This algorithm also runs in *O*(|*H*|*N* log |*H*|*N*) time. The reduction presented in Lemma 7 can be easily extended for Problem 5 as well.

## 4 Implementation and benchmarking details

### Implementation details

We implemented Algorithm 1 and the modifications to include gap penalty in Minichain v1.3 (https://github.com/at-cg/minichain). Our new chaining implementation replaces the haplotype-agnostic chaining implementation proposed in our previous work [6]. Minichain supports an end-to-end seed-chain-extend workflow to align long reads and contigs to acyclic pangenome graphs. Similar to the previous version of the software, we use the seeding and base-to-base alignment routines from Minigraph [26]. We leave the implementation of our *O*(|*Q*| | *E*| |ℋ|) time base-to-base alignment algorithm (Section 2.3) as future work because it requires further practical optimizations to reduce the runtime, e.g., using banded alignment or wavefront heuristics [37,47,64].

In Minichain, we parse the set of vertices, edges, and haplotype paths from a graphical fragment assembly (GFA) file [21]. A read may be sampled from the same or the opposite orientation with respect to the graph. Accordingly, for each connected component in the input graph, we create another component of the same size with reverse-complemented vertex labels and reversed edges. We use the same path cover for the second component as the original component but the direction of each path is reversed. We compute anchors by using Minigraph’s seeding method, which finds (*w, k*)-minimizer matches [51] between the query sequence *Q* and the vertex string labels *σ*(*v*) for all *v* ∈ *V*. The default values of *w* and *k* are 11 and 17 respectively. We set the weight of each anchor as 200 *· k* by default based on our experimental observations. Thus for *k* = 17, each anchor in a chain contributes 3400 to the total chain score. Minichain can report multiple best scoring chains and assign a mapping quality [29] to each chain; the criteria for this remains same as before [6].

### Pangenome graph construction

We used 60 publicly-available complete major histocompatibility complex (MHC) haplotype sequences [27]. This set of 60 MHC haplotype sequences comprises the CHM13 MHC haplotype [41] and 59 other haplotypes from the Human Pangenome Reference Consortium [31]. The length of these sequences varies from 4.8 Mbp to 5.1 Mbp. The MHC region is known to have significant polymorphism, inter-gene sequence similarity and long-range haplotype structures [8]. We used Minigraph (v0.20-r559) [28] to build an acyclic pangenome graph using the 60 MHC haplotypes. The graph construction algorithm in Minigraph augments structural variants (SVs) of size ≥50 bp in the graph. Our MHC pangenome graph contained 984 vertices, 1, 387 edges, and 210 SVs. After computing the graph, we realigned each haplotype to the graph to obtain the corresponding haplotype paths.

### Simulation of query sequences

We simulated 135 MHC haplotype sequences. Each sequence was simulated as an imperfect mosaic of the 60 reference haplotypes (Figure 5). We used the following procedure for simulation: (i) Select a random haplotype path in the graph, (ii) Read the first *l* bases from the chosen haplotype path in the graph, (iii) If the (*l* + 1)^*th*^ base is in vertex *v* and |*haps*(*v*)| ≥2, then switch to another randomly chosen haplotype path at vertex *v*, (iv) Read the next *l* bases, and repeat the procedure until we hit the last base of a haplotype path. After generating a sequence, we added substitutions at randomly chosen loci in the sequence. These substitutions approximately mimic true mutations and sequencing errors in real data. For each substitution rate of 0.1%, 1% and 5%, we simulated 45 sequences. In each case, three sets of 15 sequences were simulated using *l* = 1 Mbp, *l* = 2 Mbp and *l* = 3 Mbp respectively. Using this procedure, the number of recombination events in all the simulated sequences ranged from 1 to 4. Our pangenome graph and the query sequences are available on Zenodo (DOI: 10.5281/zenodo.10663934).

**Fig. 5:**
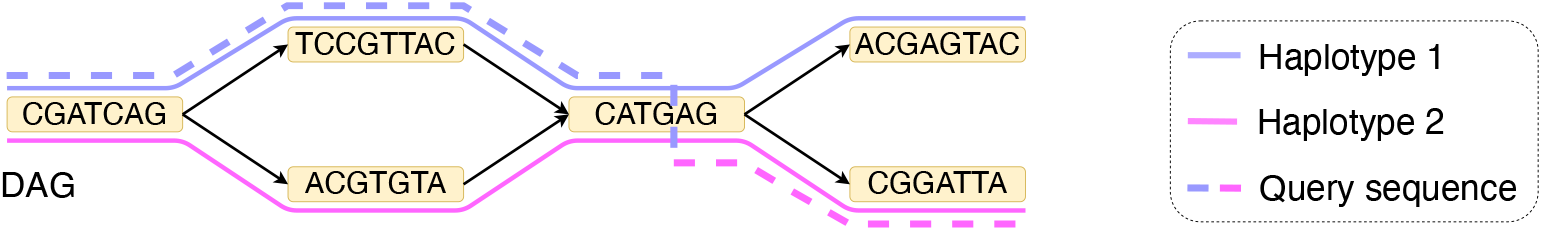
An example to illustrate simulation of a query sequence as a mosaic of reference haplotypes. In this example, *l* = 19.

## 5 Experimental results

### Comparison with haplotype-agnostic chaining using simulated data

We demonstrate that integrating recombination penalty in the sequence-to-graph chaining problem formulation leads to an improved concordance of the haplotypes used in our chains and the ground-truth. A positive finite recombination penalty in our problem formulation allows the user to limit the recombinations in an optimal chain. If we set recombination penalty *γ* = 0, then our algorithm is equivalent to the haplotype-agnostic chaining algorithm in [6]. Recombination penalty *γ* = ∞ corresponds to haplotype-restricted chaining, where a chain can use only one of the reference haplotypes. We evaluated Minichain by using simulated MHC haplotype sequences and different values of recombination penalty *γ* = 0, 10^3^, 10^4^, 10^5^, 10^6^, ∞. Minichain computes an optimal chain as a sequence of pairs of the selected anchors and the haplotype indices. Thus, we know the sequence of haplotype indices and the number of recombinations in each chain.

We show the Pearson correlation coefficients between the observed number of recombinations and the true number of recombinations using different values of substitution rates and recombination penalties (Figure 6). The results suggest that *γ* = 10^4^ gives the best correlation across different substitution rates. The correlation is weak when *γ* = 0 or *γ* = 10^6^. The correlation is not defined when *γ* = ∞ because the observed number of recombinations remained zero; thus the standard deviation of the observed values remained zero. *γ* = 10^3^ is too low for setting recombination penalty because a single anchor in a chain contributes 3400 to the chaining score (Section 4). The algorithm may still favor an additional anchor even if it involves a haplotype switch.

**Fig. 6:**
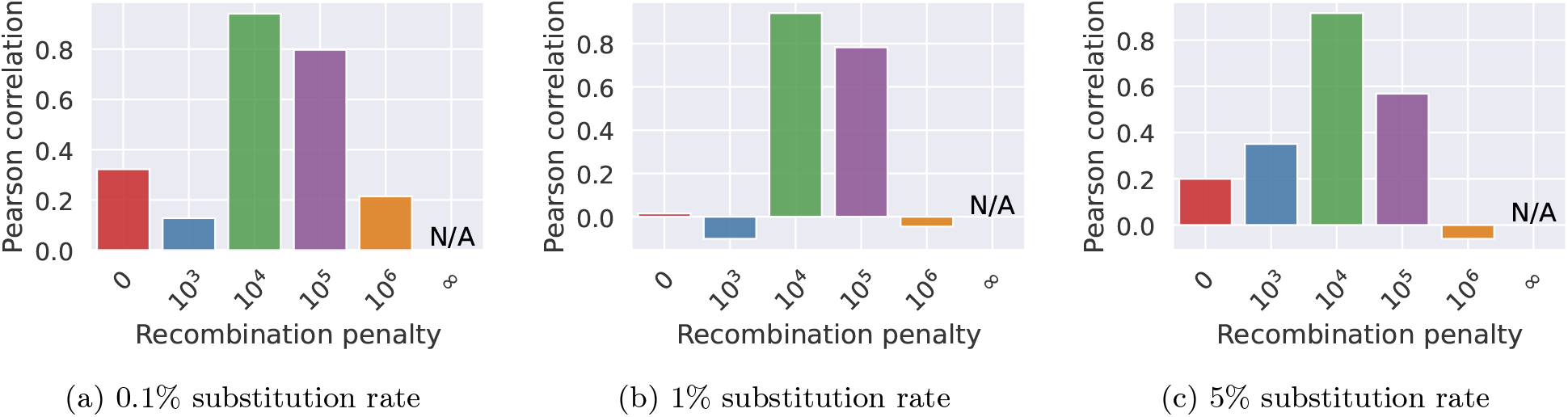
Pearson correlation between the number of recombinations in Minichain’s output chain and the true count. We evaluated the performance by using different substitution rates and recombination penalties.

Next, we compared the list of query sequences’ source haplotypes to the selected haplotypes in the output chains. Suppose a query sequence was simulated using haplotypes in the order 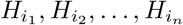, then we recorded the “true haplotype recombination pairs” as a set of pairs 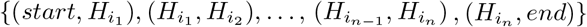. Similarly, we recorded the sets corresponding to the “observed haplotype recombination pairs” from Minichain’s output chains. We compared these two sets for each query and calculated the F1-scores (Figure 7 and Supplementary Table 1). We achieved higher F1-scores with *γ* = 10^4^, 10^5^ compared to the haplotype-agnostic (*γ* = 0) and haplotype-restricted (*γ* = ∞) modes. These results suggest that haplotype-aware chaining and alignment algorithms can be useful to prevent spurious recombinations.

**Fig. 7:**
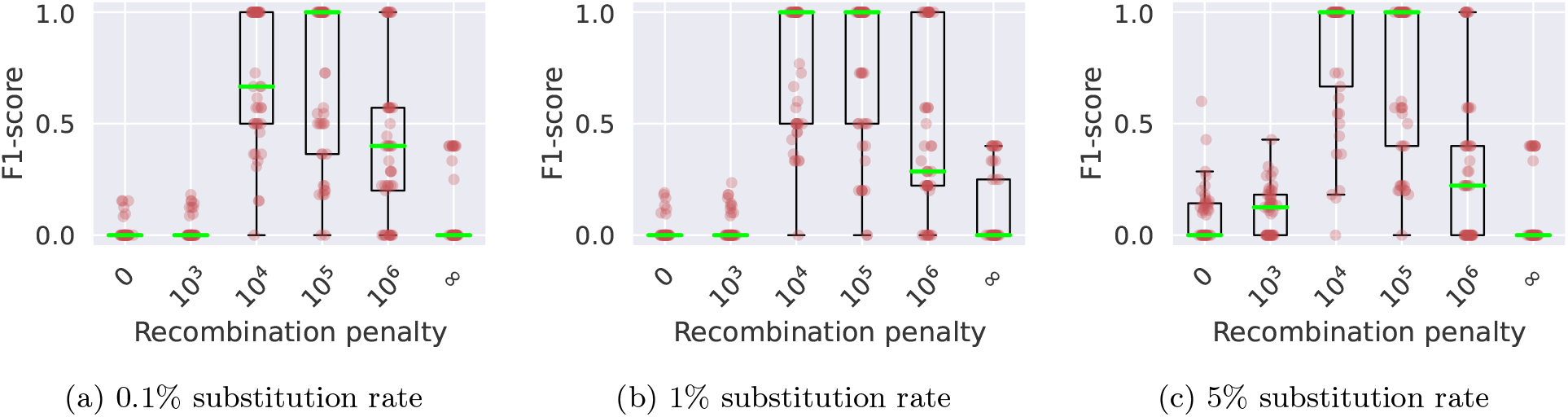
Box plots showing the levels of consistency between the haplotype recombination pairs in Minichain’s output chain and the ground-truth using three different sets of simulated MHC sequences with substitution rates (a) 0.1%, (b) 1%, and (c) 5%. We tested using different recombination penalties. Each red dot in the plots corresponds to a query sequence. The median values are highlighted in green.

Our MHC pangenome graph comprises SVs only. We expect further improvements in the accuracy of our algorithm if substitution and short indel variation are also augmented in a pangenome graph [19]. This is because more variants will help to distinguish between near-identical or closely-related haplotypes. Doing this will also require improvements in Minichain’s seeding and chaining implementation to allow the use of anchors that cover small bubbles in a graph. This is possible by considering a more flexible definition of anchors that can span multiple vertices [50,49,52].

The runtime of Minichain ranged from 5 to 11 minutes for aligning a simulated MHC sequence to the pangenome graph. The worst-case time complexity of the proposed haplotype-aware chaining algorithm is higher compared to the existing haplotype-agnostic chaining algorithms. This is because we require haplotype paths as the path cover, whereas the haplotype-agnostic chaining algorithms use a path cover of minimum size [6,32,35]. For example, our graph with 60 haplotype paths has a minimum path cover of size 5. Thus, we observed that our haplotype-aware chaining algorithm in Minichain was about 10*×* slower compared to the haplotype-agnostic chaining algorithm [6]. Space requirements of our algorithm are also proportional to the count of haplotype paths (e.g., to store the score table *C*). As a result, we observed that the haplotype-aware algorithm required about 15*×* more memory compared to the haplotype-agnostic algorithm [6].

### Evaluation using real data

We evaluated Minichain by using real long-read sequencing datasets from CHM13 human genome. Our MHC pangenome graph includes CHM13 as one of the 60 haplotypes. Since the CHM13 genome is effectively haploid, we expect all chains to be consistent with the CHM13 haplotype path in the graph. We used two PacBio HiFi datasets (SRX5633451, SRR11292121) and one Oxford Nanopore dataset (SRR23365080). We filtered the reads sampled from MHC region by mapping the reads to T2T-CHM13v2.0 genome reference (GCF_009914755.1) using minimap2 v2.26 [26]. Next, we discarded reads of length ≤ 1 kbp. After these filtering steps, the N50 read lengths of the three datasets were 11 kbp, 20 kbp, and 83 kbp, respectively. We considered the chaining output of a read as correct if all the anchors in the highest scoring chain overlapped with the CHM13 haplotype path in the pangenome graph. The fraction of correctly chained reads using the haplotype-agnostic objective (*γ* = 0) on the three datasets were 97.3%, 96.1%, and 91.1%, respectively. Using *γ* = 10^5^ as recombination penalty, the fraction improved to 97.8%, 97.2%, and 94.2%, respectively. These results, although preliminary, suggest that the proposed haplotype-aware sequence-to-graph alignment algorithms can be useful for genotyping and variant calling using pangenome references.

## 6 Discussion

Pangenome graph has emerged as a powerful alternative representation as a reference during resequencing of a genome. These graphs can be represented in either a haplotype-agnostic or haplotype-aware manner. The former excludes the information of the reference haplotypes, whereas the latter incorporates it. Leveraging the haplotype path information during read-to-graph or contig-to-graph mapping can be useful to compute accurate alignments and avoid unlikely haplotype recombinations. This will become even more important in near future as more reference-quality human genomes are produced and included in a human pangenome graph. Prior work on haplotype-aware mapping has focused on restricting the recombination count to either zero or one [38,57,56].

In this paper, we showed how the routine chaining and alignment tasks can be rigorously formulated and solved in a haplotype-aware manner. Our formulations focus on optimizing the standard metrics, e.g., edit distance for alignment and chaining score for co-linear chaining, while considering the switches used between the reference haplotypes. Thus, the proposed algorithms will help address the combinatorial challenges as more variants are included in a graph. The other key theoretical contribution is that we proved that our alignment and chaining algorithms are near-optimal assuming SETH. Subsequently, we demonstrated the practical value of our haplotype-aware chaining algorithm. We developed a proof-of-concept implementation in Minichain (v1.3). Our experiments show that the algorithm produces alignment with fewer false recombinations and better selection of the correct haplotypes.

More work is needed, especially to establish that the proposed haplotype-aware mapping algorithms can lead to improvements in routine genomic applications. This requires a more careful engineering of Minichain to make it useful for graphs that comprise SNPs and indels. The current implementation of Minichain is restricted to acyclic pangenome graphs that contain structural variation only. It will also be interesting to investigate whether the genotype imputation accuracy can be improved using the proposed algorithms on low-pass whole-genome sequencing data.

## Acknowledgements

This work is supported by funding from the National Supercomputing Mission, India under DST/NSM/ R&D_HPC_Applications, the Science and Engineering Research Board (SERB) under SRG/2021/000044, and the Intel India Research Fellowship. We used computing resources at the C-DAC National PARAM Supercomputing Facility, India and the National Energy Research Scientific Computing Center, USA.

## Supplementary Document

### Algorithm for solving Problem 5

To efficiently compute gap costs between anchors during co-linear chaining, we follow the idea as described in [6]. We assume the same definition of gap cost function as used in the Problem 2a of reference [6]. First, we precompute an index of the DAG to facilitate efficient calculation of gaps during chaining. Let *D*(*v*_1_, *v*_2_) denote the shortest path measured in terms of the number of characters from *v*_1_ to *v*_2_. If path *P*_*h*_ covers vertex *v*, then suppose *dist*2*begin*(*v, h*) is the number of characters in haplotype *h* before vertex *v*. Accordingly, our precomputed index contains *last*2*reach*(*v, h*), *D*(*last*2*reach*(*v, h*), *v*) and *dist*2*begin*(*v, h*) for all *v* ∈ *V* and *h* ∈ [|ℋ|]. All these quantities can be computed in *O*(|*E*| |ℋ|) time during preprocessing of the DAG [6]. See Algorithm 2 for an outline of the gap-sensitive haplotype-aware chaining procedure. This is similar to Algorithm 1 except that we introduce two variables *g*_1_ and *g*_2_ (Lines 18, 27) to consider gaps between anchors. The arguments in the proofs of Lemma 6 in this paper and Lemma 4 in reference [6] can be easily extended to argue the correctness of Algorithm 2.

#### Algorithm 2

*O*(|*ℋ*|*N* log |*ℋ*|*N*) time chaining algorithm for Problem 5

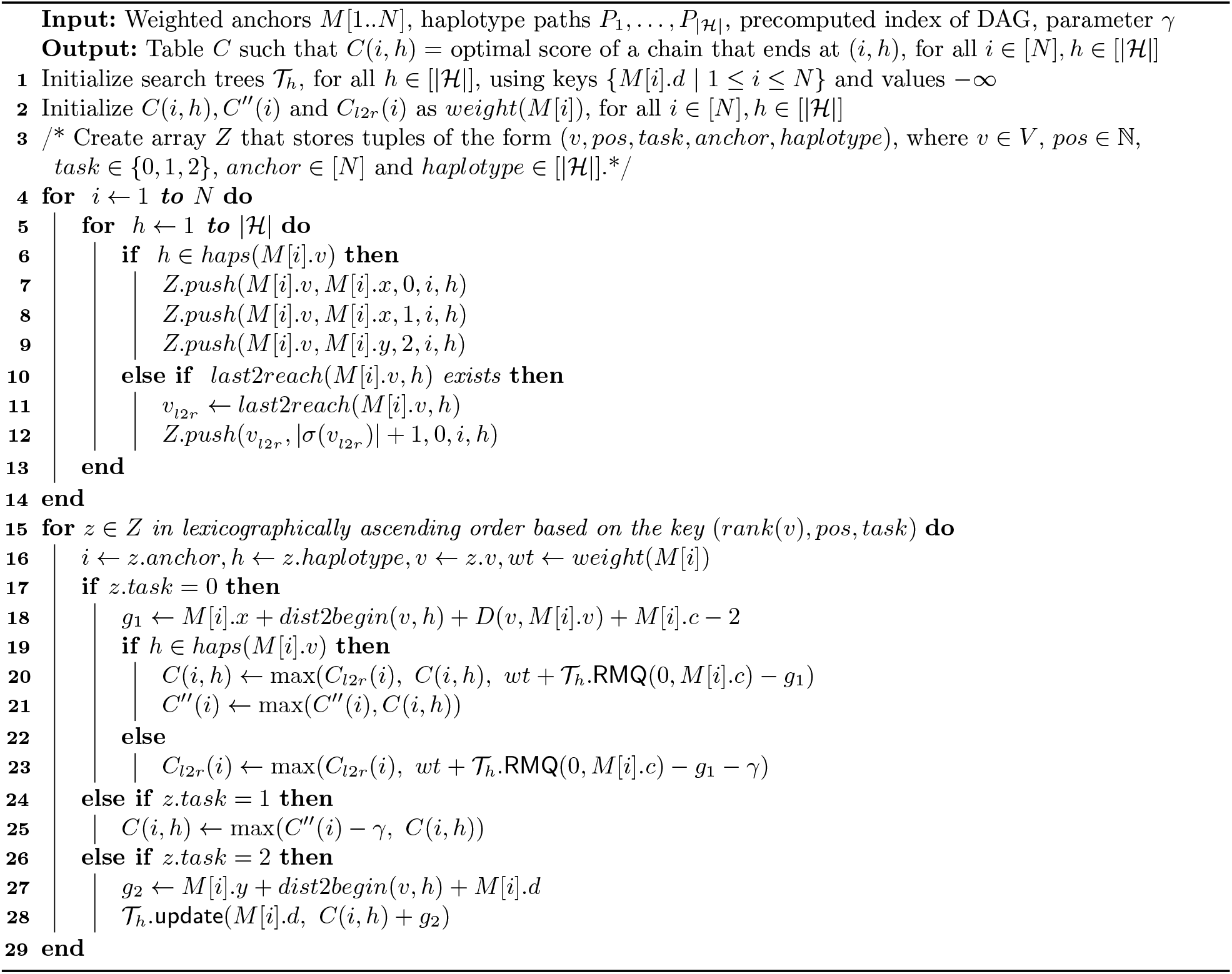

**Table 1:**
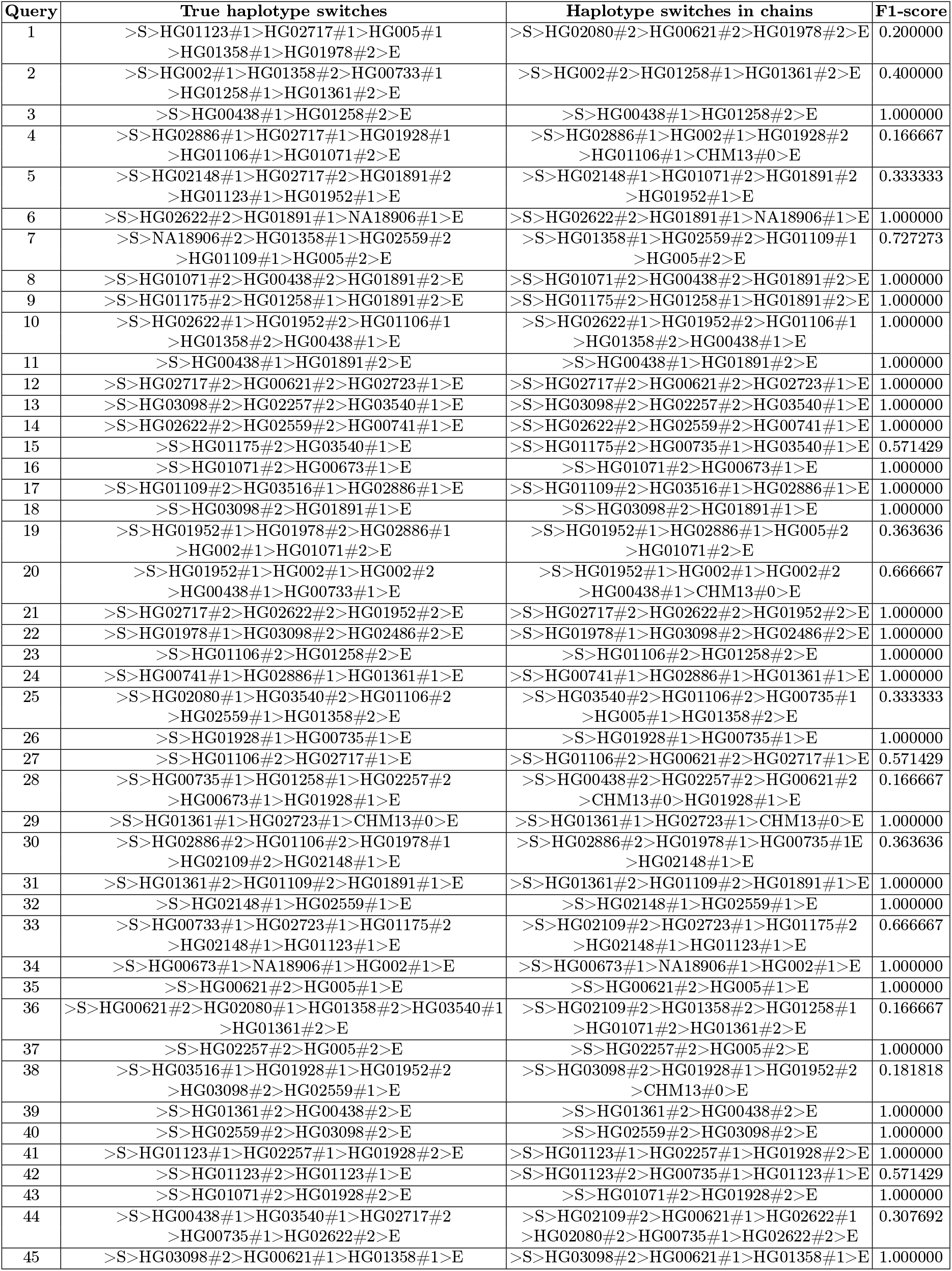
We show the true recombinations between haplotypes and the recombinations observed in co-linear chains using recombination penalty *γ* = 10^4^. We show this data for 45 query sequences that were generated using 0.1% substitution rate. ‘S’ indicates the start and ‘E’ indicates the end.

